# Confidence as a diagnostic tool for perceptual aftereffects

**DOI:** 10.1101/270280

**Authors:** Regan M. Gallagher, Thomas Suddendorf, Derek H. Arnold

**Author notes:** Corresponding author: Regan Gallagher.

## Abstract

Perceptual judgements are, by nature, a product of both sensation and the cognitive processes responsible for interpreting and reporting subjective experiences. Changed perceptual judgements may thus result from changes in how the world appears (perception), or subsequent interpretation (cognition). This ambiguity has led to persistent debates about how to interpret changes in decision-making, and if cognition can change how the world looks, or sounds, or feels. Here we introduce an approach that can help resolve these ambiguities. In three motion-direction experiments, we measured perceptual judgements and subjective confidence. Sensory encoding changes (i.e. the motion-direction aftereffect) impacted each measure equally, as the perceptual evidence informing both responses had changed. However, decision changes dissociated from reports of subjective uncertainty when non-perceptual effects changed decision-making. Our findings show that subjective confidence can provide important information about the cause of aftereffects, and can help inform us about the organisation of the mind.

---

A central challenge in perception research is to understand how the world looks, feels, and sounds, as opposed to how it is remembered, imagined, or judged. But perceptual judgements are, by nature, a product of both sensation and the cognitive operations responsible for producing measurable behaviour. Changes in perceptual decision making could thus equally arise from changes to sensory encoding and perception, or from later decisional processes that operate independently of perception. This ambiguity has fostered persistent debate regarding the degree to which our cognitions — like imaginations, motivations, or beliefs — can change the sensory processes that determine what we perceive^1,2,3,4,5^. Here we introduce a new approach that can help shed light on whether or not changes in perceptual judgements reflect changes in sensation and perception.

Much of what we know about human perception has resulted from investigating sensory aftereffects^6,7^. An aftereffect is a change in the measured boundary between perceptual categories, which can result from prolonged and repeated exposures to a specific stimulus^8^, or rapidly from a single brief exposure to a test stimulus^9,10,11^. It remains a matter of debate, however, how best to dissociate perceptual from post-perceptual effects on decision making^12,13^. We examine this problem by using motion-direction judgements and subjective confidence, and we show that reports of high and low confidence provide important information about the cause of changes in perceptual decisions.

Perceptual decisions are often measured by forced-choice categorisations. Participants might be tasked with determining which binary category a test stimulus belongs, for example motion direction (left or right), orientation (clockwise or counter-clockwise tilt), or facial characteristics (masculine or feminine). Stimuli that appear ambiguous (e.g. incoherent motion, near vertical orientations, or androgynous faces) represent the boundary between these categories and, in an unbiased observer, are equally likely to be categorised as belonging to either category.

Metacognition research shows that an individual can accurately predict their own ability to discriminate between perceptual categories^14,15^. Decisions likely to be correct carry greater feelings of confidence, whereas decisions likely to be incorrect (or made by guessing) carry lower confidence. The individual’s boundary between categories is their point of subjective equality (PSE), characterised by both probabilistic responses and subjective uncertainty.

Since typical observers can accurately rate their own performance in perceptual decision tasks, confidence might provide important information about whether an aftereffect represents a change in a perceptual or cognitive boundary. On one hand, aftereffects caused by changes to sensory encoding should equally impact categorical decisions and confidence reports, because the sensory evidence underlying both judgements has changed. On the other hand, aftereffects arising only from changes to decision processes^12,13^ might selectively or disproportionately impact decisions made under uncertainty, without corresponding changes in confidence (see Figure 1).

**Figure 1:**
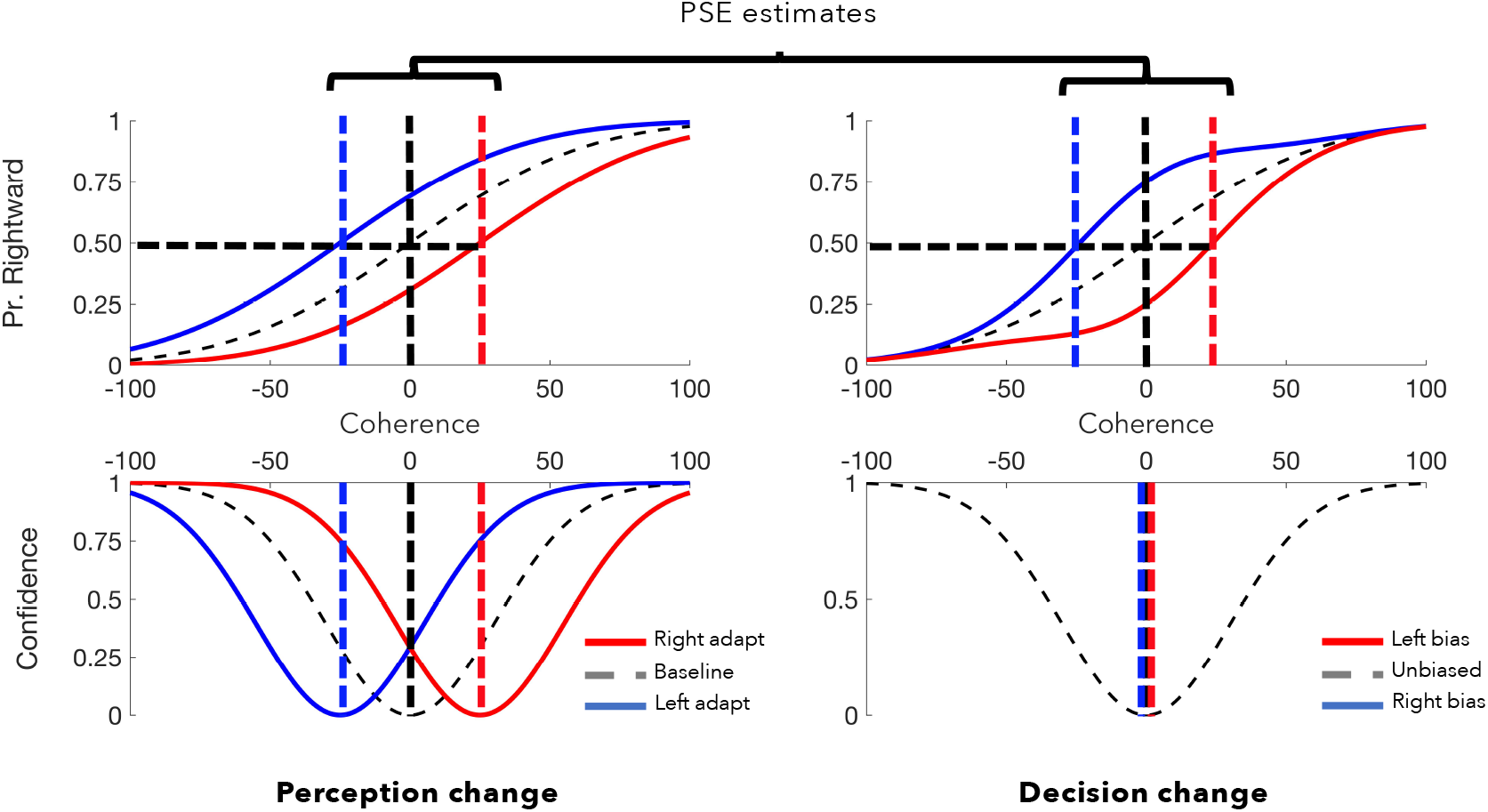
Confidence can distinguish perceptual from non-perceptual effects on decision making. Here we illustrate hypothetical data sets for motion-direction judgments. Categorical decisions (“are stimulus elements predominantly moving left, or right?”) are plotted above as a function of motion coherence (the proportion of elements physically moving left or right). Expressions of confidence (“how confident are you that your decision is correct?”), as a function of motion coherence, are plotted below. Confidence is assumed to scale with the strength of sensory evidence. On the left we depict patterns of results expected from changed sensory evidence, on the right we depict patterns expected from people making different decisions about ambiguous inputs. If sensory evidence changes, previously ambiguous inputs should *look* as if they are moving right (blue line) or left (red), so categorical decisions and confidence are expected to shift in tandem as both judgments are informed by sensory evidence. If, however, decisions change because people make different decisions regarding ambiguous inputs, categorical decisions might shift without any change in expressions of confidence.

It might seem odd to suggest that people could make different perceptual decisions even if perception is unchanged. This is possible, however, if people are implicitly or explicitly encouraged to make different decisions about ambiguous information. An instruction to dwell on immoral actions, for instance, could implicitly encourage people to regard ambient lighting as ‘darker’ than if they dwelt on moral actions^4,16^. Or people might have a tendency to repeat decisions that were effective in the past, which could result in ambiguous inputs being judged as more similar to previous inputs^17^. Either scenario could result in people making different decisions about ambiguous information, even if perception is unchanged. Crucially, as neither scenario involves changes to perception, the range of inputs associated with lowest confidence should also be unchanged. If this is true, confidence could be used to resolve ambiguity about the underlying cause of many aftereffects. To test this, we measured decisions and associated confidence in three motion-direction experiments.

In Experiment 1 we examined the motion-direction aftereffect, where decision changes are known to reflect changes to physiological processes^18^. As predicted, we find that categorical direction decisions and confidence measures are equally impacted by adaptation. In Experiment 2 we instruct participants to default to biased decision strategies when they encounter ambiguous tests. This experiment demonstrates that categorical decisions can change without any change to perception, and when this happens, expressions of confidence remain unchanged. Finally, in Experiment 3, we examine serial dependency (the impact of prior tests on subsequent judgments). Results suggest that the most recent stimulus (1-back) changes perception, impacting categorical decisions and confidence equally. The stimulus before last (2-back), however, seems to *selectively* impact categorical decisions, consistent with a post-perceptual influence on decisions. Overall, our data show that confidence reports can provide diagnostic evidence for determining whether or not changes in decision-making reflect changes in perception.

## Experiment 1

We first checked that aftereffects can be equally estimated from categorical decisions and confidence judgements.

People made categorical direction decisions (left / right) and confidence judgments (low / high) about tests that varied in direction and motion coherence. Cumulative Gaussian functions were fit to each participant’s distribution of rightward direction judgements as a function of motion coherence (see Methods). The 50% points were taken as estimates of the point of subjective equality (PSE) — the stimulus value equally likely to be judged as moving left or right. A raised Gaussian function was fit to each participant’s distribution of low-confidence responses, and the peak of the fitted function was taken as a second PSE estimate the point of peak uncertainty. All *t*-tests reported are two-tailed repeated measures tests for equality of means. All Bayes’ factors were estimated using JASP software^19^, with the default Cauchy prior width of 0.707.

### Baseline

To illustrate our approach, in Figure 2 we depict distributions of categorical decisions (top) and expressions of uncertainty (or low-confidence; bottom). As can be seen, functions fit to these distributions can provide a closely matched PSE estimate (direction decisions, M = −0.43, SD = 1.60; confidence reports, M = −0.51, SD = 1.03; difference *t*_14_ = 0.18, *p* = .862, BF_10_ = 0.27). These baseline data are averaged across all participants, but PSE estimates were derived from individual function fits.

**Figure 2:**
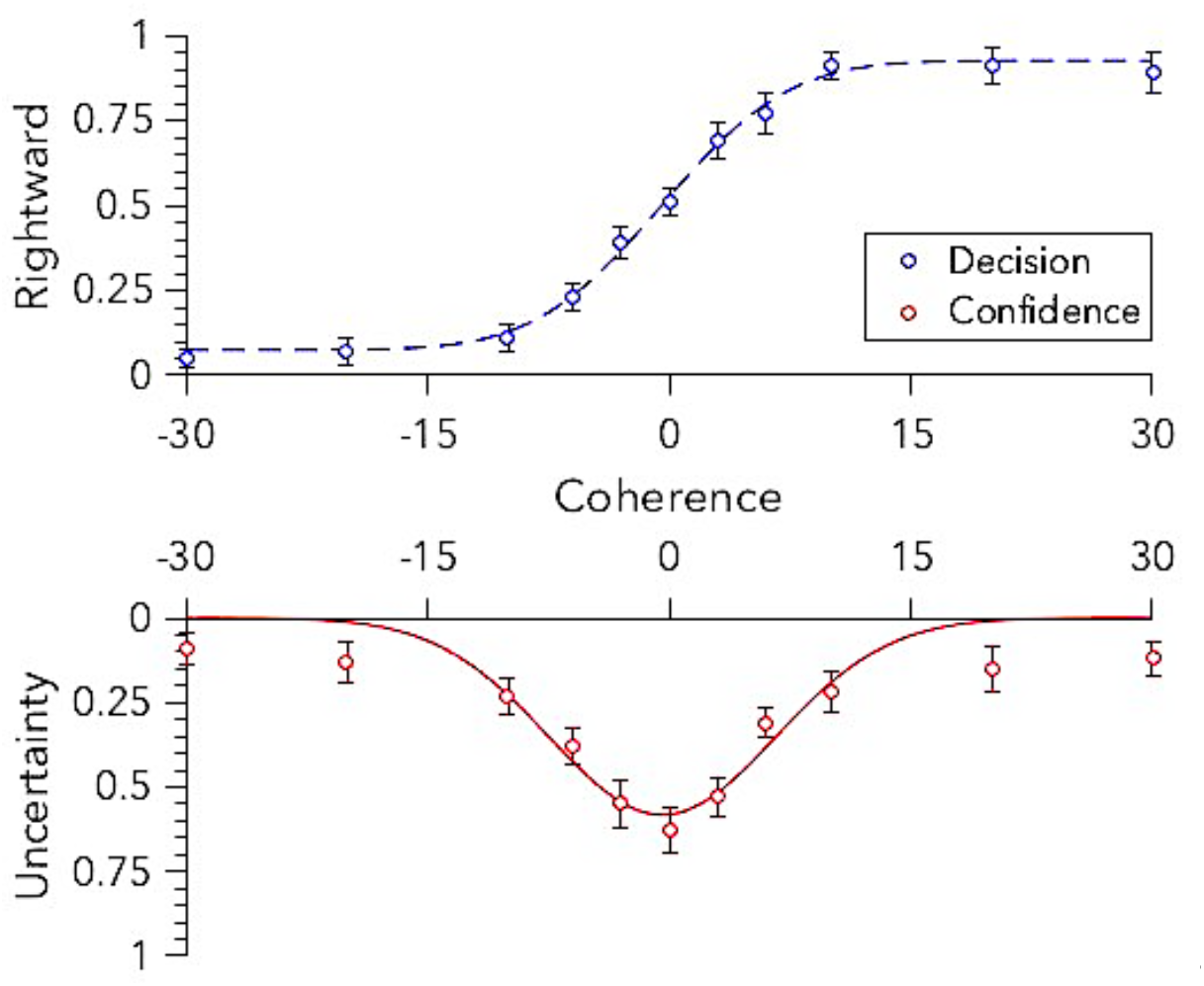
Baseline data (N=15). Distributions of reported motion direction (left or right) and confidence (high or low) as a function of dot coherence (percent) and direction (negative values left, positive right). Data and best-fit functions are depicted for baseline motion-direction judgments in Experiment 1. The inflection point of the function fitted to decision data (blue), and the peak of the function fitted to confidence data (red) each estimate the point of subjective equality. Depicted data are averaged across all participants, but PSE estimates are derived from individual fits. Error bars depict ±1 SEM.

### Coherent motion adaptation

In the adaptation phase, participants adapted to a dot motion stimulus with 30% coherence, moving either to the left or right (see methods for further details). Adaptation data for each participant produced two decision distributions, each indicating the proportion of stimuli judged as having moved predominantly rightward as a function of test dot coherence and direction. One distribution depicts responses for left-adapted trials, the other for right-adapted trials. The impact of adaptation was estimated as the difference between these two PSE estimates for each individual. Adaptation data for each participant also produced two distributions of confidence, indicating the proportion of low-confidence responses as a function of test dot coherence and direction (again for left-adapted and right-adapted trials).

Adaptation to coherent motion had a robust impact on direction decisions, with left adaptation (L_PSE_ = −11.67, SD = 7.31) and right adaptation (R_PSE_ = 11.22, SD = 6.76) producing different PSE estimates (*t*_14_ = 6.90, *p* < .001). These data are depicted in Figure 3 (left panel, top). The same pattern of results was observed for confidence data. Adaptation robustly impacted measures of uncertainty, with left adaptation (L_CONF_ = −10.02, SD = 5.79) and right adaptation (R_CONF_ = 9.93, SD = 5.27) producing different confidence profiles (*t*_14_ = 7.98, *p* < .001). The effect of adaptation on decision data (ΔPSE = 22.89, SD = 12.84) and confidence data (ΔCONF = 19.95, SD = 9.69) was not statistically different (*t*_14_ = 1.73, *p* = .106). These data are also depicted in Figure 3 (left panel, bottom).

**Figure 3:**
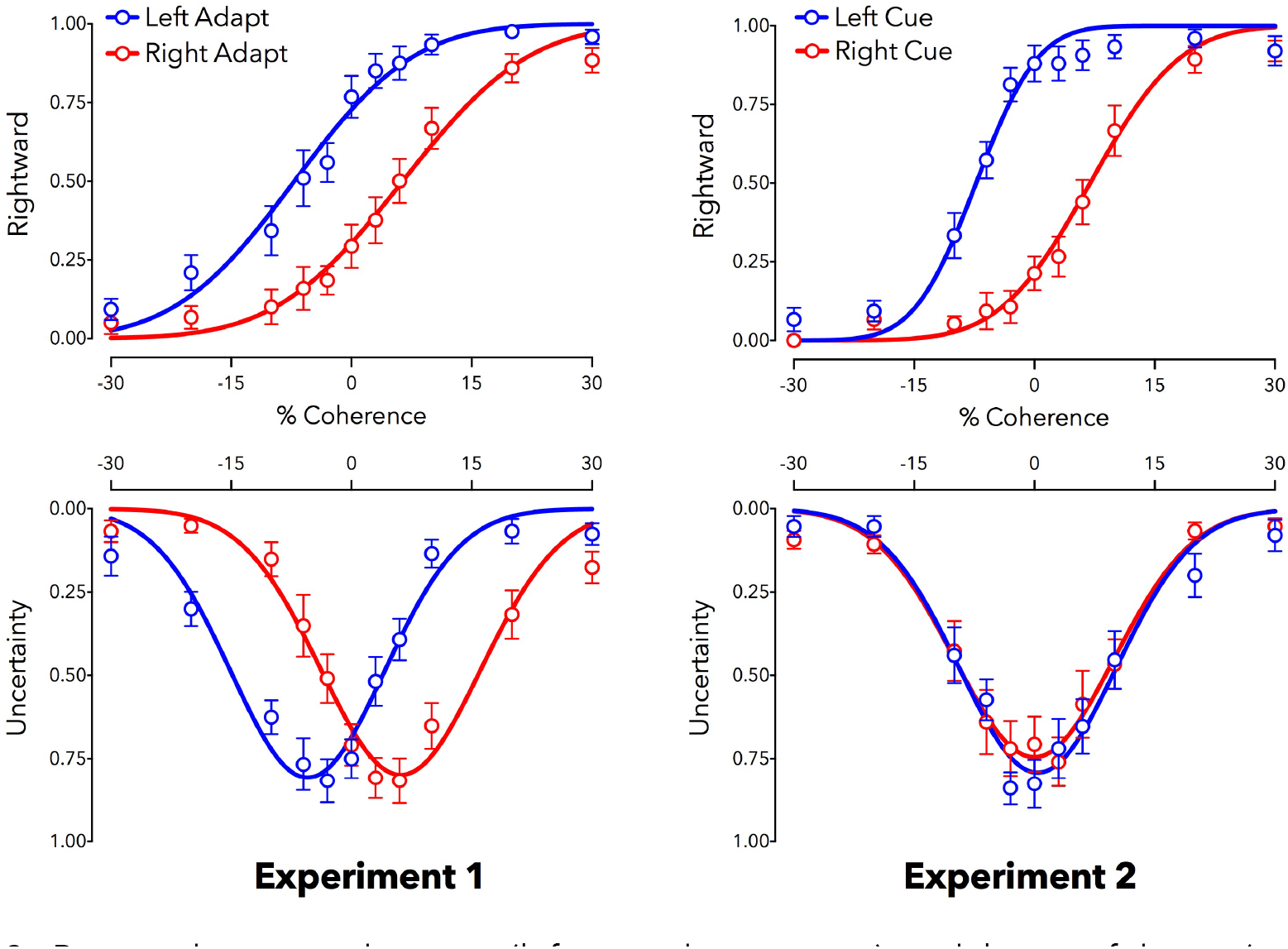
Reported motion direction (left or right, top row) and low-confidence (uncertainty, bottom row) as a function of dot coherence (percent) and direction (negative values left, positive right). Data are shown along with best-fit functions for adaptation data in Experiments 1 (left column) and 2 (right column). Both Experiments are associated with changed direction decisions (see top row), but only Experiment 2 is associated with shifts in confidence (bottom row; see main text for further details). Depicted data are averaged across participants, but statistical tests were based on individual data fits. N = 15; Error bars depict ±1 SEM.

The results of null hypothesis statistical tests are also supported by Bayes Factor analyses, with extremely strong evidence favouring adaptation-induced changes (decision BF_10_ = 2.92 x 103; confidence BF_10_ = 1.28 x 104). Cohen’s *d* effect size estimates for each measure show that adaptation produces extremely large aftereffects (decisions *d* = 1.78, 95% CI 0.94 – 2.60; confidence *d* = 2.06, 95% CI 1.14 – 2.96). It is important to note that decision and confidence measures provide a statistically equivalent measure of the aftereffect (note the overlapping CIs). This is important as it argues against future dissociations arising because confidence measures provide a less sensitive measure of adaptation. The two measures have provided an equivalent estimate of an aftereffect known to result from physiological changes that impact perception^18,20^.

## Experiment 2

Experiment 1 showed that the motion aftereffect can be measured equally well using categorical perceptual decisions and confidence judgments. In Experiment 2 we assess the logical counterpoint: an aftereffect that results from people making different decisions about ambiguous inputs, without any perceptual changes. To achieve this, in Experiment 2 people ‘adapt’ to motion with no coherent direction. Participants were then instructed to make a default decision (either left or right, according to instruction) when they encounter subjectively ambiguous tests. This experiment tested the robustness of each response method against decision biases.

### Adapt to random motion

Participants ‘adapted’ to a random dot motion stimulus (0% coherence). This should have no systematic impact on sensory encodings of motion^20^. A static direction cue, an arrow pointing to the left or right, was also presented above the ‘adaptor’ (see Methods for further details). We instructed participants to adopt a default decision, rather than guess, if they were unable to determine the direction of a test. This prompted people to report either the direction consistent (congruent condition) or inconsistent (incongruent condition) with the arrow when test motion direction was ambiguous. The congruent condition is expected to produce category decisions consistent with an assimilative aftereffect (when tests are judged as being *similar* to an ‘adaptor’)^21^. The incongruent condition is expected to produce categorical decisions changes consistent with a negative aftereffect (when tests are judged as being *dissimilar* to an ‘adaptor’, as in the classic motion-direction aftereffect measured in Experiment 1). The important question is the degree to which confidence judgments are impacted by systematically biased categorical decisions about ambiguous inputs.

#### Categorical direction decisions

When faced with an ambiguous test, participants were more likely to report that the test stimulus was moving in the same direction as the cue in the congruent task, and this bias impacted PSE estimates (L_PSE_ = 8.21, SD = 6.58; R_PSE_ = −7.15, SD = 3.48; difference, *t*_14_ = 7.50, *p* < .001). Participants were less likely to report the same direction in the incongruent task, which also impacted PSE estimates (L_PSE_ = - 8.81, SD = 4.44; R_PSE_ = 7.77, SD = 4.40; difference, *t*_14_ = 8.33, *p* < .001; see Figure 3). These were extremely large effects (congruent *d* = 1.93, 95% CI 1.05 – 2.79; incongruent *d* = 2.15, 95% CI 1.20 – 3.08). Bayes Factor analyses additionally favoured the hypotheses that decision biases can produce data consistent with both assimilative (congruent BF_10_ = 6.62 x 103) and contrastive (incongruent BF_10_ = 2.01 x 10^4^) aftereffects (see Figure 3).

#### Confidence results

When faced with an ambiguous test, participants were no more likely to report low (or high) confidence for different tests in either the congruent (L_CONF_ = −0.12, SD = 1.13; R_CONF_ = 0.62, SD = 1.07; difference, *t*_14_ = 1.15, *p* = .268) or incongruent (L_CONF_ = 0.02, SD = 1.26; R_CONF_ = −0.38, SD = 1.02; difference, *t*_14_ = 0.85, *p* = .410; see Figure 3) tasks.

#### Dissociation of decision and confidence effects

There was a clear dissociation between PSE estimates from direction decisions and confidence judgments in the congruent (decision ΔPSE = 13.46, SD = 6.20; confidence ΔCONF = 0.50, SD = 1.68; difference, *t*_14_ = 9.23, *p* < .001) and incongruent (decision ΔPSE = 14.94, SD = 7.91; confidence ΔCONF = 0.41, SD = 1.85; difference, *t*_14_ = 7.07, *p* < .001) tasks. Effect size estimates for these differences were extremely large (congruent *d* = 2.01, 95% CI 1.11 – 2.89; incongruent *d* = 1.62, 95% CI 0.83 – 2.39). The dissociation is supported by Bayes Factor analyses, which provide extremely strong evidence for the alternate hypothesis, that categorical decision and confidence measures differed (congruent BF_10_ = 6.14 x 10^4^; incongruent BF_10_ = 3.71 x 10^3^).

In combination, Experiments 1 & 2 show that categorical decisions and confidence judgements can equally measure an aftereffect known to result from physiological processes that change perception (Experiment 1), but provide dissociable measures when ‘aftereffects’ instead result from people making different decisions about ambiguous inputs (Experiment 2). Having validated our proposal in these opposite contexts, we adopted it to assess the likely cause of serial dependence.

## Experiment 3

Recent studies have established a contingency between perceptual decisions and preceding tests, without any need for protracted adaptation periods^9,10^. One interpretation is that serial dependence results from rapid sensory adaptation, which measurably impacts perception on a trial by trial basis. Another interpretation suggests that serial dependence can result from post-perceptual aspects of decision making, like working memory^11^, or from people repeating their previous response when subsequent stimuli are ambiguous^17^. We assessed these proposals by measuring serial dependencies between sequential categorical decisions and confidence judgments.

People made categorical direction decisions (left / right) and confidence judgments (low / high) about tests that varied in direction and motion coherence, akin to baseline trials in Experiment 1 (see Methods for further details).

### N-back effects

Trial responses were subdivided according to the direction of the last test (1-back data), and according to the direction of the test two trials prior (2-back data). Functions were fit to these data to quantify 1-back and 2-back effects on categorical decision making, as a function of motion direction and coherence on the present test (see Methods for further details).

#### 1-back effects

Participants tended to categorise tests as moving in the *opposite* direction relative to the last test (L_PSE_ = −1.50, SD = 2.38; R_PSE_ = 1.72, SD = 3.18; difference, *t*_21_ = - 4.46, *p* < .001; see Figure 4, left panel, top). The last trial also impacted confidence judgments. Peak uncertainty shifted in tandem with direction judgments, in the *opposite* direction relative to the last test (L_CONF_ = −1.99, SD = 1.74; R_CONF_ = 1.35, SD = 1.64; difference, *t*_21_ = −7.46, *p* < .001; see Figure 4).

**Figure 4:**
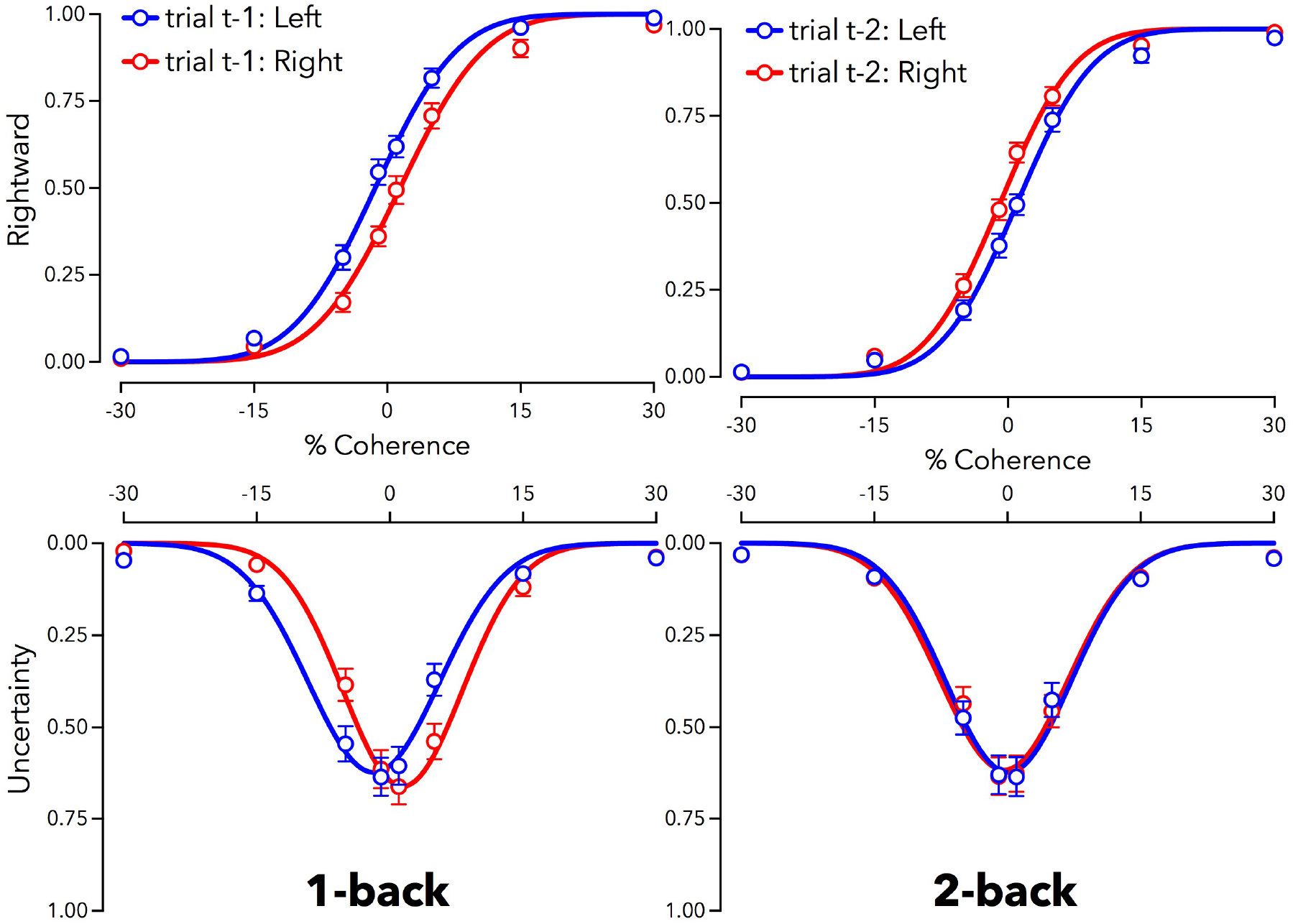
Proportion of rightward motion reported (top row) and low-confidence (uncertainty, bottom row) as a function of dot coherence (percent) and direction (negative values left, positive right). Data are divided according to the physical direction of the last test (Left panel: t-1 data), and according to the physical direction of the test two trials ago (Right panel: t-2 data). Blue data indicate that the prior trial was moving left, red that the prior trial was moving right. Depicted data are averaged across participants, but statistical tests were based on individual fits. N=22; Error bars depict ±1 SEM.

1-back effects can be described as large, both for categorical decisions (decision *d* = 0.95, 95% CI 0.44 – 1.45) and confidence (*d* = 1.59, 95% CI 0.95 – 2.22). Again, it is important to note that confidence provided a statistically equivalent measure of this serial dependency relative to categorical decisions (note the overlapping CIs). Bayes Factor analyses provided strong support for the hypothesis that the last trial impacted direction decisions (decision BF_10_ = 1.39 x 10^2^) and confidence judgments (BF_10_ = 6.52 x 10^4^). These effects were statistically equivalent (decision ΔPSE = 3.23, SD = 3.39; confidence ΔCONF = 3.34, SD = 2.10; difference, *t*_21_ = 0.20, *p* = .841), and a Bayes Factor analysis provided moderate evidence for the equivalence of the two 1-back effects (decision vs confidence BF_10_ = 0.23). This pattern of results is consistent with those obtained for the classic motion-direction aftereffect in Experiment 1, suggesting that serial dependence effects can be driven by processes that impact perception.

#### 2-back effects

Participants tended to categorise tests as moving in the *same* direction as the test two trials prior (t-2; L_PSE_ = 1.34, SD = 2.98; R_PSE_ = −1.20, SD = 2.45; difference, *t*_21_ = 3.44, *p* = .002; see Figure 4). There was, however, no discernible impact of this trial on confidence judgements (L_CONF_ = 0.09, SD = 1.37; R_CONF_ = −0.48, SD = 1.87; difference, *t*_21_ = 1.45, *p* = .161). There was also a significant dissociation between measures of aftereffect estimates (categorical decision ΔPSE = 2.54, SD = 3.46; confidence ΔCONF = 0.58, SD = 1.87; difference, *t*_21_ = 2.81, *p* = .011). These data are consistent with the pattern of results in Experiment 2, where an ‘aftereffect’ resulted from people making different decisions about ambiguous inputs, rather than from sensory encoding changes.

The effect size estimate for 2-back categorical decision effects was moderately large, as was the effect size for differences between the decision and confidence judgments (decision *d* = 0.73 95% CI 0.26 – 1.20; decision / confidence difference *d* = 0.60 95% CI 0.14 – 1.05). Bayes factor analyses provided moderate to strong evidence in favour of both the decision effect and the dissociation between decision and confidence judgments (decision BF_10_ = 16.51; decision vs confidence difference BF_10_ = 4.78). Figure 5 compares changes in categorical decisions and confidence judgments across all three experiments.

**Figure 5.**
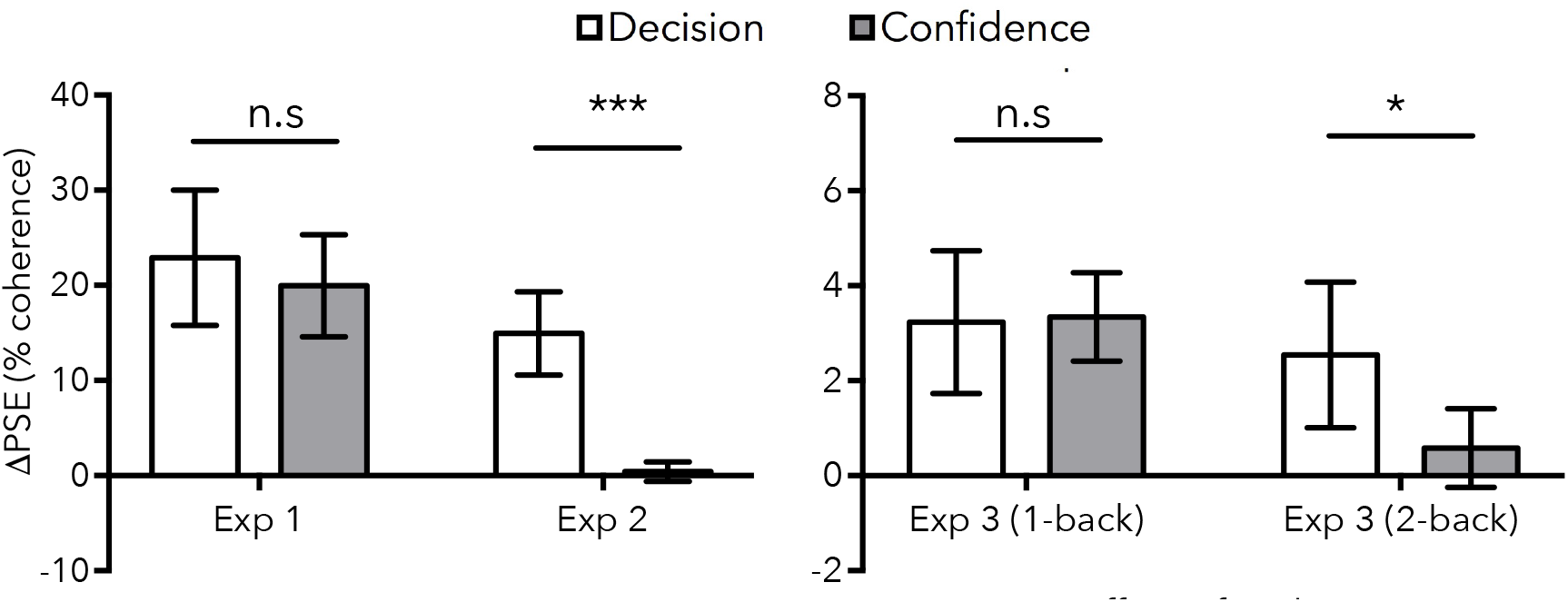
Bar graphs summarising the results of all experiments. Data depict mean (absolute) PSE changes calculated from categorical decisions (white bars) and confidence judgments (grey bars). Error bars depict 95% CIs.

## General Discussion

Our data indicate that confidence provides important diagnostic information for distinguishing sensory encoding aftereffects from aftereffects that do not change perception. Experiment 1 showed that an aftereffect driven by sensory adaptation^18,20^ can be measured equally well using categorical decisions and confidence judgments (see Figures 2 & 3). Experiment 2 showed that a non-perceptual decision bias (an instruction to make systematically biased decisions about ambiguous inputs) had different impacts on categorical decisions and confidence; whereas categorical decisions provided evidence consistent with large aftereffects, confidence judgments suggested no change in perception (see Figure 3). This implies that confidence can be robust against decision biases that favour one category response over another when sensory input is ambiguous.

Having validated our approach in opposite contexts, Experiment 3 examined serial dependencies between successive moving tests. Our results suggest that sensory adaptation can indeed occur rapidly^9,10^. We observed an equal impact of the last trial on subsequent categorical decisions and confidence judgments (see Figure 4, left, and Figure 5, 1-back results). These data mirror our results for an aftereffect known to be caused by sensory adaptation (Experiment 1), so these results suggest that the last test can act as an adapting stimulus, generating a contrastive aftereffect that rapidly changes perception.

Our Experiment 3 data also reveal serial dependence had an assimilative influence on decision making^10,21^. The stimulus two trials prior to a test (2-back) had an *attractive* impact on categorical decisions. However, there was no evidence for a 2-back assimilative aftereffect for confidence judgments (see Figure 4, right, and Figure 5, 2-back results). These results mirror the results of Experiment 2, where people were instructed to make systematically biased categorical decisions when inputs were ambiguous. The 2-back effect in Experiment 3 is likely driven by people being biased to repeat categorical decisions when inputs are ambiguous^17^, consistent with a post-perceptual influence of decision making on perceptual judgements^11^.

Our study shows that aftereffects estimated exclusively from categorical decisions only provide ambiguous evidence in favour of perceptual changes. Here we have shown that this evidence should be augmented with confidence measures. Using the classic motion-direction aftereffect^18,20^ (see Experiment 1), we show that confidence is equally impacted by physiological processes that change perception. However, in Experiment 2, we showed that confidence measures could remain veridical when categorical decisions change — these data are otherwise suggestive of an ‘aftereffect’, but arose because people were making systematically biased judgments about perceptually ambiguous tests. These data suggest that adding confidence judgments to categorical decision protocols (which are typically *exclusively* used to measure perceptual aftereffects) can reveal additional diagnostic information about the likely cause of a measured aftereffect.

It is important to note that confidence was a highly sensitive measure for the aftereffects that arose because of physiological processes that change perception (see Figure 6, compare Experiment 1, & Experiment 3: 1-back results). When dissociations between categorical decision and confidence measures arose (compare Experiment 2, & Experiment 3: 2-back results), it is unlikely that this could be attributed to confidence judgments having provided a poor, or insensitive, measure of a perceptual aftereffect.

**Figure 6.**
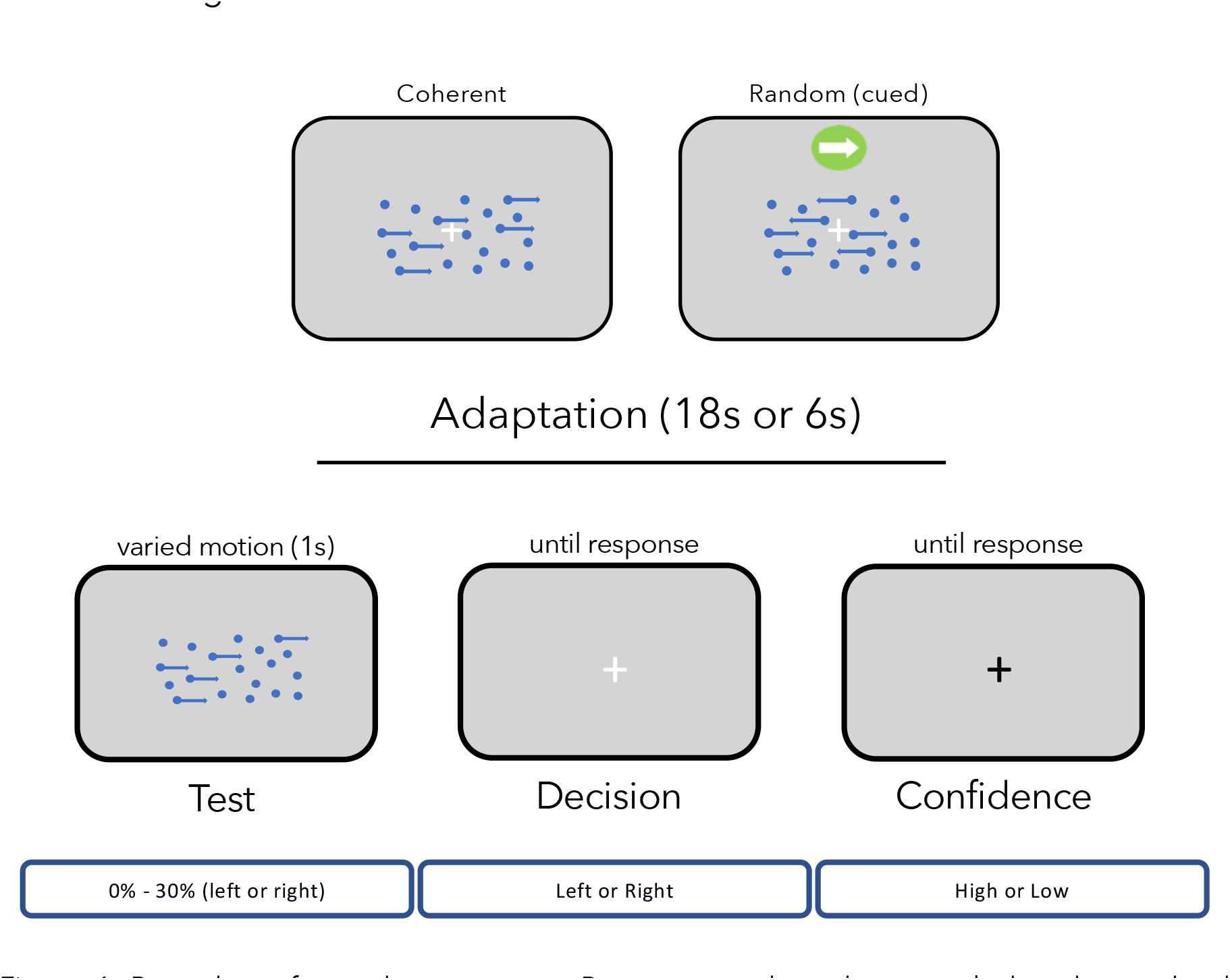
Procedures for each experiment. Participants adapted to stimuli that depicted either coherent motion or random motion and a static direction cue. Each trial within a testing block consisted of an adaptation phase (except for baseline) followed by a dot test probe. The direction of the adapting stimulus (left or right) was consistent within the first half of a block, and then changed direction for the second half. Adapting stimuli appeared for 18s on the first trial of each block, and on the middle trial when the stimulus changed direction, and for 6s on all other trials. Dot test probes were present for 1s, appearing on the second frame after the adapting stimulus disappeared. A new trial began once participants had recorded their direction decision (left or right) and reported their confidence (whether they had confidence that their decision was correct—yes / high confidence or no / low confidence).

Our Experiments demonstrate that a participant’s insight into their own decision making can resolve many contemporary debates in perception. Confidence reports leverage this insight to provide additional diagnostic information about the likely cause of an aftereffect. When decision changes result from changes to perception, the evidence informing both perceptual judgements and associated confidence has changed. In this case, an aftereffect should equally impact both measurements. However, when an aftereffect arises because people are making different decisions about perceptually ambiguous inputs, perceptual decisions should dissociate from reported confidence. In this case, aftereffect measures selectively impact decision making under uncertainty, and are likely not indicative of a perceptual aftereffect.

Many studies have measured the effect of adaptation to high-level stimuli^2,3,4,21,22,23^, and have observed robust changes in categorical perceptual decisions. However, the extent to which aftereffects of this kind represent top-down changes to perception remains in dispute^1,3,12,13,16,23^. Expanding upon our results might help resolve current ambiguities about the causes of high-level aftereffects, and provide a means of directly testing for top-down effects of cognition on perception^1,3,5^.

Experiments examining sensory aftereffects have been vital to developing our understanding of the physiological processes that underlie perception and perceptual decision making. Ultimately, however, we would like to be able to differentiate perceptual and post-perceptual processes, because people can make different decisions even when inputs might look, feel, or sound identical. This dilemma can be addressed by identifying the range of inputs that elicit greatest uncertainty — which can be achieved by asking participants to report on their confidence in each categorical perceptual decision.

## Conclusion

Confidence reports can provide additional information for determining if an aftereffect changes how the world looks, or sounds, or feels. Our approach could augment traditional protocols, at little cost in time and effort.

## Method

### Participants

All participants were recruited from the University of Queensland’s Psychology department, and were naïve as to the purpose of the experiments. Experiment 1 had a sample size of 15, which provides very high power to detect perceptual adaptation effects (these are typically large; Cohen’s d > 0.8). This conservative sample size estimate was chosen because the effect of adaptation on confidence was yet unknown. Experiment 2 also had a sample size of 15 to match Experiment 1. The sample size for Experiment 3 was increased to 24 because serial dependence effects are smaller than adaptation effects (although still moderately large, Cohen’s d > 0.5). Two participants were excluded from Experiment 3 because either their category judgements (n=2) or confidence judgements (n=1) could not be adequately modelled using a standard psychometric function fit, so the final sample size of Experiment 3 was 22. Participants in Experiment 2 were postgraduate Psychology and UQ Perception Lab members, all other participants were recruited from a first-year student pool.

### Ethics

Ethical approval for all experiments was obtained from the University of Queensland’s (UQ) Ethics Committee, and all experimental tasks were performed in accordance with the UQ guidelines and regulations for research involving human participants. Each participant provided informed written consent to participate in the study and were made aware that they could withdraw at any moment from the study without prejudice or penalty. First year students received course credit in exchange for participation.

### Stimuli and apparatus

In all tasks the motion stimuli were generated using Matlab software and the Psychophysics Toolbox^24,25^. Stimuli were presented on Dell P791 monitors (1024×768 pixels) with a refresh rate of 60Hz. The dot-motion stimuli were single-pixel dots rendered blue (RGB = 0,0,255). Coherent motion signals were created by translating dots in the coherent direction by 1 pixel on each frame. To avoid participants tracking individual dots, a new subset of dots (equal to the % dot coherence) were selected at random on each frame. All other dots were moved to a new and randomly selected location on each frame, within the aperture, to create a background of incoherent motion.

Test stimuli in all experiments consisted of 100 blue dots, shown against a grey background. Dot coherence values ranged from −30 (30% leftward) through 0 (random motion) to +30 (30% rightward). Test stimuli were set to one of 11 coherence values [−30 −20 −10 −6 −3 0 3 6 10 20 30], presented in a random order.

In Experiment 1, adapting stimuli consisted of 100 dot stimuli, each 1 pixel in size. A coherent motion signal was achieved by selecting 30 dots at random on each frame to be displaced left or right by one pixel. All other dots were redrawn at random locations. Again, to avoid participants tracking individual dots, a new subset of dots (equal to the % dot coherence) were selected at random on each frame. All other dots were moved to a new and randomly selected location on each frame, within the aperture, to create a background of incoherent motion. Adapting motion was in one direction for the first half of the block of trials, and in the other direction for the second half of the block, with the initial adaptation direction determined at random for each participant.

In Experiment 2 ‘adapting’ stimuli were similar to Experiment 1, with two exceptions. First, motion coherence was 0, so *all* dots were redrawn at random locations on each frame. Second, a static direction cue was presented directly above the stimulus window — an arrow pointing to the left or right (see Figure 6). The static direction cue was in one consistent direction for the first half of a block of trials, and reversed for the second half of the testing block. The initial cue direction was determined at random for each participant.

In Experiments 1 and 2, each test stimulus was presented 10 times, and repeated for each adaptation or cue direction (totalling 220 stimulus observations). There were no adapting stimuli in Experiment 3, so each test stimulus was viewed 50 times (totalling 550 stimulus observations).

### Procedure

Participants received a written instruction of the experimental procedure. After reading the instructions, participants were then verbally queried whether they understood how to report their responses. All understood the direction judgement was to probe which direction the test appeared to move (left or right), and how to respond if they were unsure (which was varied according to experiments). Likewise, all participants understood to report whether they had high or low confidence in each response or, if it was more intuitive, to report whether they thought their guess was likely correct or incorrect. All participants verbally acknowledged that they understood both the direction and the confidence responses.

When participants had acknowledged they understood the instructions, they then sat comfortably in front of the monitor at a distance of approximately 55 cm, rested their fingers on the keyboard’s arrow buttons, and fixated a central cross-hair. If there was an adaptation phase, the adapting stimulus was presented for 18 seconds on the first trial in each block, and again on the middle trial (when adapting motion direction reversed). On all other trials the adapting stimulus lasted for six seconds.

On each test trial, participants were presented with one of the 11 tests for 1 second. After the stimulus presentation period concluded, participants could immediately indicate whether the test had appeared to be moving left (by pressing the left arrow key) or right (by pressing the right arrow key). In Experiments 1 and 3, participants were instructed to provide their best guess if they could not determine which direction the test was moving. In the adaptation (but not baseline) phase of Experiment 2, participants were instructed to default their response to the direction congruent (Experiment 2a) or incongruent (Experiment 2b) with the static reference cue presented on that trial. Participants in Experiment 2 completed both tasks (congruent and incongruent) in a random and counterbalanced order.

Once participants had completed a direction judgment, the white fixation cross hair turned black, which prompted the participant to make a confidence judgment. Participants could immediately report whether they felt their response was probably correct (by pressing the up arrow – a high confidence response) or probably wrong (by pressing the down arrow – a low confidence response). The fixation cross turned white once the confidence response had been recorded, and a new trial started after a 50ms delay.

## Additional information

### Authorship Declaration

R. M. Gallagher and D. H. Arnold developed the concept and experimental designs. R. M. Gallagher tested participants and performed statistical analyses. R. M. Gallagher drafted the manuscript under the supervision of T. Suddendorf, & D. H. Arnold, who provided critical revisions. All authors wrote and approved the final version of the manuscript.

